# Persistence of backtracking by human RNA polymerase II

**DOI:** 10.1101/2023.12.13.571520

**Authors:** Kevin B Yang, Aviram Rasouly, Vitaly Epshtein, Criseyda Martinez, Thao Nguyen, Ilya Shamovsky, Evgeny Nudler

## Abstract

RNA polymerase II (pol II) can backtrack during transcription elongation, exposing the 3’ end of nascent RNA. Nascent RNA sequencing can approximate the location of backtracking events that are quickly resolved; however, the extent and genome wide distribution of more persistent backtracking is unknown. Consequently, we developed a novel method to directly sequence the extruded, “backtracked” 3’ RNA. Our data shows that pol II slides backwards more than 20 nucleotides in human cells and can persist in this backtracked state. Persistent backtracking mainly occurs where pol II pauses near promoters and intron-exon junctions, and is enriched in genes involved in translation, replication, and development, where gene expression is decreased if these events are unresolved. Histone genes are highly prone to persistent backtracking, and the resolution of such events is likely required for timely expression during cell division. These results demonstrate that persistent backtracking has the potential to affect diverse gene expression programs.

## Introduction

RNA polymerase II (pol II) produces all transcripts for protein coding genes and many noncoding RNAs in eukaryotes. At certain positions, pol II can backtrack along the template, disengaging the 3’ end of nascent RNA from the catalytic site and extruding it through the pore and funnel domain (secondary channel) resulting in transcription pausing or long-term arrest^1–3^. Productive elongation can be promptly restored by transcription factor IIS (TFIIS), which promotes cleavage of the extruded, “backtracked” RNA, thereby generating the new RNA 3’ terminus in the catalytic site of pol II^4–7^.

Backtracking in both eukaryotes and bacteria has been implicated in gene regulation and DNA repair ^1,8–13^. TFIIS can stimulate transcription of genes in response to heat shock and hypoxia^14–16^. In addition, unresolved backtracking can be deleterious to cells due to interference with transcription^17^ and transcription-replication collisions causing DNA breaks in bacteria^18,19^ and yeast^9^.

Genome wide characteristics of backtracking has previously been inferred in eukaryotes by comparing shifts in pol II pauses in wild-type and null or cleavage deficient TFIIS cells^14,20^. Specifically, the position of pol II pauses shifts downstream since the 3’ end of backtracked RNA is un-cleaved. However, this measures cleavage events and not backtracking itself; therefore, it is only sensitive to transient events that are quickly resolved *in vivo*. Persistent backtracking events that are more slowly resolved would be undetectable using these comparisons. Extensive *in vitro* characterization of long-ranged backtracking has shown that it is slower to be resolved because the backtracked RNA binds a site within the secondary channel that stabilizes the complex and impedes TFIIS binding^2^. Contrarily, shorter backtracking can be rapidly resolved by either intrinsic or TFIIS cleavage^21^.

In this work, we systematically map and functionally characterize persistent backtracking events in mammalian cells with **Lo**ng **Ra**nge Cleavage (LORAX) sequencing, a novel method that directly sequences backtracked RNA to precisely determine where backtracking begins and ends with nucleotide resolution *in vivo*. Our data shows that persistent backtracking primarily occurs where pol II pauses, however pause strength does not determine the potential for persistent backtracking. Using crosslinked RNA footprinting of pol II, we show that pol II can be captured in a stable position proximal to the nascent 3’ end of RNA at pauses with persistent backtracking. We show that persistent backtracking occurs predominantly near promoters and intron-exon junctions. Moreover, it is not stochastic and has a propensity to occur in certain genes, namely those involved in translation, cell cycle, nucleosome assembly and development. In cells where persistent backtracking cannot be resolved, the expression of such genes is decreased. Histone genes are highly prone to persistent backtracking, and we provide evidence that the accumulation and resolution of these events is critical for the timely expression of histones during replication. From this data we conclude that persistent backtracking has the potential to affect diverse gene expression programs.

## Results

### Detection of transient backtracking in mammalian cells

Previous efforts to map genome-wide backtracking have used nascent transcription sequencing to profile cells unable to cleave backtracked RNA^14,20^. As a result, the 3’ end of nascent RNA extends downstream, and a corresponding shift in mNET-seq peaks occur between wild-type and TFIIS null mutants, allowing for the indirect detection of backtracking events (Figure 1A).

**Figure 1.**
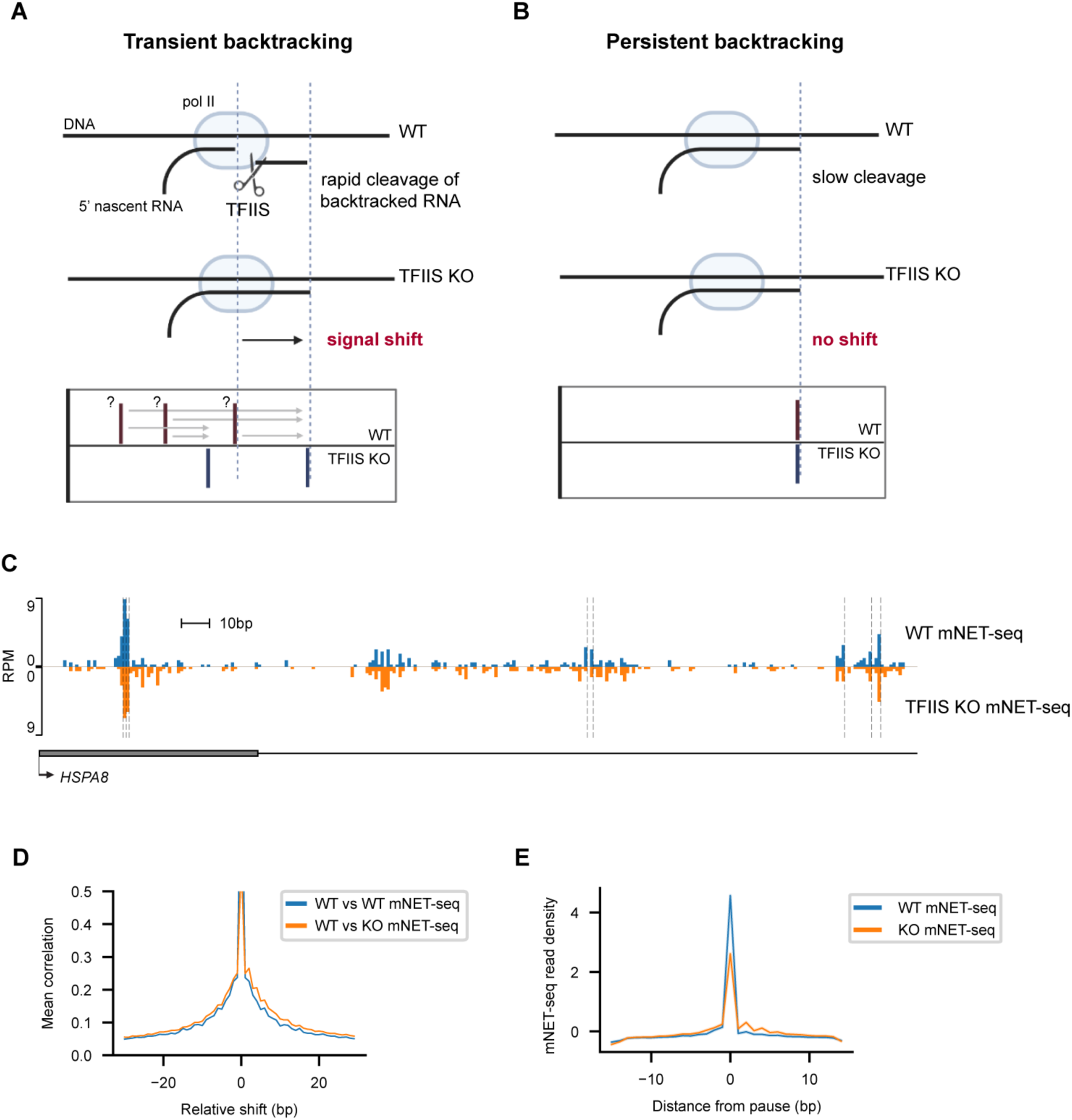
Detection of transient backtracking in mammalian cells. (A) Schematic of transient backtracking detected by comparison of nascent transcription profiles of wild-type (WT) and TFIIS knockout (KO) cells. TFIIS KO prevents cleavage of backtracked RNA causing a downstream shift in signal. The appearance of this shift requires that cleavage occurs rapidly in wild-type cells, and that cleaved complexes can be captured. (B) Schematic of persistent backtracking which cannot be detected by nascent transcription sequencing comparisons. (C) Representative genome viewer of mNET-seq signals derived from WT (top) and TFIIS KO (bottom) cells across the promoter of HSPA8. Pauses are denoted by dotted vertical lines. (D) Mean cross correlation between WT and KO mNET-seq data of well transcribed genes (top n=1,000 genes sorted by RPKM) was calculated by determining the Pearson’s correlation between fixed KO and shifted WT data, averaging over all genes, and compared to WT cross correlation. (E) Meta-analysis of WT and TFIIS KO mNET-seq signal around the strongest n=100,000 pauses found in WT mNET-seq. Reads within 15 bp of WT mNET-seq pauses were aggregated, normalized within each window, and then averaged. All mNET-seq experiments were conducted in duplicate for each condition.’

Importantly, this approach is limited to detecting unstable, transient backtracking that is quickly resolved by wild-type TFIIS *in vivo* since it detects backtracked complexes after cleavage before transcription resumes, and not the backtracking event itself (Figures 1A and S1A). This approach is also unable to determine the length and precise position where backtracking begins and ends because it is impossible to distinguish which mNET-seq peaks correspond with the beginning or end of a specific backtracking event (Figure 1A; bottom). In the case that a backtracking event is more persistent *in vivo*, backtracked pol II would be expected to enter a post-cleavage state more rarely such that no mNET-seq shift would occur (Figures 1B and S1A). Persistent backtracking events would therefore be undetectable using mNET-seq comparisons.

Nonetheless, this approach works well for mapping transient backtracking. A large downstream shift of 5-18 base pairs (bp) can be observed in 75% of NET-seq pauses across the genome in *Δdst1* yeast cells, demonstrating that transient backtracking is ubiquitous and is rapidly resolved in yeast^20^. Interestingly, a smaller shift of less than 5 bp is observed in mNET-seq pauses prepared from mammalian cells overexpressing a cleavage-deficient, dominant negative (DN) mutant of TFIIS, possibly because the DN mutant is unable to completely prevent cleavage^14^.

To test this approach in mammalian cells completely lacking wild-type TFIIS, we generated a knockout of TCEA1 and confirmed genetic editing by sequencing and western blot of cytosolic and nuclear fractions of wild-type (WT) and TFIIS knockout (KO) cells (Figure S1B-C). We also constructed a plasmid for overexpression of TFIIS to demonstrate the specificity of the TCEA1 antibody and for subsequent experiments (Figure S1C). We then prepared mNET-seq libraries from WT and TFIIS KO cells and determined the location of pol II pausing as previously described (Figure 1C, Methods)^20,22^.

In a representative comparison of mNET-seq data for WT and TFIIS KO cells at the promoter of HSPA8, we found shifts in pol II pauses to be unpronounced (Figure 1C; pauses indicated by vertical dotted-lines). In cross correlation analysis across well-expressed genes, only a small rightward shift in signal is observed (Figures 1D and S1D-F). A genome wide meta-analysis of TFIIS KO mNET-seq signal centered around strong WT pauses also indicates a downstream shift of less than 5bp (Figures 1E and S1G-H), in agreement with what was observed with DN TFIIS (Sheridan et al., 2019). In addition, we calculate a ‘backtracking index’ by determining the difference in read density upstream and downstream of each pause in WT and KO mNET-seq as previously described^14^. By comparing the distributions of differences between conditions, we determine that only 5.8% of pol II pauses can be confidently associated with transient backtracking (Figure S1I). We also find that TFIIS KO has few effects on nascent transcription profiles, as meta-analyses around genomic landmarks such as splice, transcription start and transcription termination sites are unchanged (Figures S1J-L).

Together, the short length and minor shifts in mNET-seq comparisons, the small fraction of transiently backtracked pauses, and the minute effects of TFIIS KO on nascent transcription profiles suggest that, in contrast to yeast^20^, backtracking in mammalian cells is either a relatively rare transient event, or that it is more persistent in a way that cannot be detected by nascent RNA sequencing.

### Nucleotide resolution mapping of persistent backtracking by LORAX-seq

To directly detect persistent backtracked RNA across the human genome we developed LORAX-seq. First, transcription complexes were isolated from human cells using an adapted mNET-seq protocol with modifications from the subsequent POINT method which greatly increases the purity of immunoprecipitated transcription complexes and foregoes MNase digestion, which might degrade backtracked RNA^22,23^. Briefly, nuclei are isolated and chromatin is solubilized using urea and detergent to allow for efficient DNase digestion. Pol II is then immunoprecipitated with magnetic beads and washed to eliminate contaminating RNA and proteins^23^. To further decrease noise, purified transcription complexes were treated with phosphatase to remove contaminating 5’ phosphates as substrates for ligation. Transcription complexes were then aliquoted, and mock treated or treated with purified recombinant human TFIIS to stimulate cleavage and elution of backtracked RNA. Notably, recombinant TFIIS was highly active on backtracked transcription complexes assembled *in vitro*, cleaving a majority of RNA (Figures S2A-B). TFIIS cleavage produces new 5’ phosphates for ligation and the eluted RNA can then be purified, ligated to adaptors in a strand specific manner, and prepared for sequencing (Figures 2A and S2C, Methods). Prepared in this way, sequenced RNA directly provides the genomic position of where backtracking begins and ends (Figure 2A, bottom).

**Figure 2.**
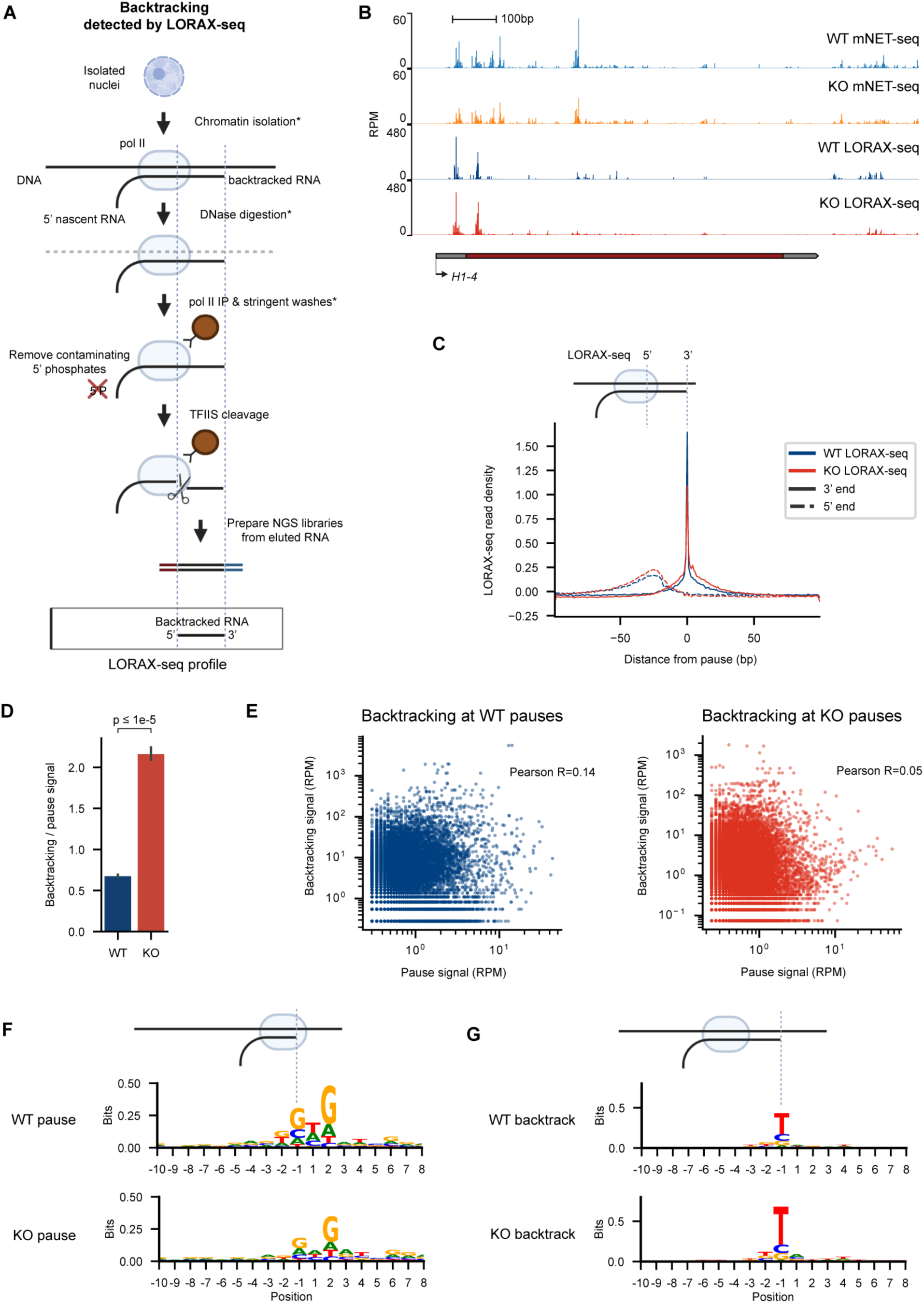
Nucleotide resolution mapping of persistent backtracking events by LORAX-seq. (A) Schematic of LORAX-seq. Chromatin was isolated from fractionated nuclei using urea and Empigen before digestion with DNase. Transcription complexes were immunoprecipitated, washed and then treated with phosphatase to remove contaminating 5’ phosphates. Purified transcription complexes were then incubated with purified TFIIS in vitro to cleave and elute backtracked RNA, generating new 5’ phosphates for ligation. The elution was purified and prepared for next generation sequencing. Sequencing reads directly correspond with backtracked RNA, providing the precise location that backtracking begins and ends. * denotes steps adapted from POINT^23^. (B) Representative genome viewer of persistent backtracking signal (LORAX-seq 3’ signal) compared to nascent transcription signal (mNET-seq) across the histone gene H1-4 from WT and TFIIS KO cells. All LORAX-seq experiments were conducted in duplicate for each condition. (C) Meta-analysis of WT and TFIIS KO LORAX-seq signal around n=100,000 pol II pauses from Figure 1E. Both the 5’ and 3’ ends of LORAX-seq is provided. Reads within 100 bp of pauses were aggregated, normalized within each window, and averaged. Schematic of where each signal is derived is provided above the plot. (D) LORAX-seq signal within 50 bp of the pause normalized to pause signal from the analysis in (C). (E) LORAX-seq signal within 50bp of the pause compared to pause signal at each pause in reads per million from WT and TFIIS KO cells from the analysis in (D). The Pearson correlation is provided. (F) Consensus sequence around n=20,000 strong pol II pauses extracted from mNET-seq from WT and TFIIS KO cells. Schematic of the signal under consideration is provided above the plot. (G) Consensus sequence around n=20,000 sites of persistent backtracking from LORAX_seq data from WT and TFIIS KO cells. Schematic of the signal under consideration is provided above the plot.

Following preparation of LORAX-seq libraries, we observed that TFIIS treatment increases elution of short RNA species from both WT and TFIIS KO transcription complexes (Figure S2C-D). After sequencing, we observed that TFIIS treated libraries had significantly more reads that were uniquely mapped and aligned to protein coding sequences when compared to mock treated libraries in biological replicates (Figures S2E-F). We also observed that more protein-coding RNA was eluted from TFIIS KO pol II which also occurred to a smaller degree in the absence of TFIIS treatment, suggesting that background signal in the mock treated samples arises from self-cleavage by pol II (Figures S2C-F). The increase in protein-coding RNA eluted from TFIIS KO transcription complexes reflect an increase in unresolved backtracking *in vivo*, indicating the specificity of the protocol for detecting backtracking *in vivo*.

After mapping, we find backtracked RNA to mainly be between 20 and 40 nucleotides in length (Figure S2G). We note that mapping to the mammalian genome limits our analysis of persistent backtracking to reads longer than 20 nucleotides. Additionally, we find LORAX-seq to be robust since backtracking events mapped to each gene are reproducible in biological replicates (Figures S2H-I). Reverse transcriptase mispriming is known to introduce artifacts during library preparation; however, these artifacts are limited to <3% of signals in both our mNET-seq and LORAX-seq datasets (Figure S2J)^24,25^.

When comparing LORAX-seq to mNET-seq data across a representative gene, we find that persistent backtracking is more prevalent at certain pol II pauses (Figure 2B). To compare persistent backtracking signal to nascent transcription genome wide, we generated meta-analyses of LORAX-seq signal centered around pol II pauses extracted from mNET-seq (Figure 1E). We find that the 3’ end of backtracking signal is closely aligned with pol II pauses, while the 5’ end is distributed between 40 and 20 bp upstream (Figures 2C and S2K-L). Interestingly, TFIIS KO LORAX-seq 5’ ends are found more consistently at the -20nt position (Figures 2C and S2K-L). This indicates that backtracking length is more variable in WT LORAX-seq data when compared to TFIIS KO (Figure S3A). Perhaps WT cells are quicker to resolve backtracking at this distance while TFIIS KO cells are not. From this analysis, we determine that persistent backtracking events generally begin at positions where pol II pauses and are between 20 to 40 bp in length.

Additionally, we observe a small downstream shift of the 3’ signal in TFIIS KO, as seen with mNET-seq, suggesting that transient and persistent backtracking events are not mutually exclusive (Figure 2C). Short and transient backtracking that is unresolved may precede longer, more persistent events.

From the same analysis, we can calculate the amount of persistent backtracking associated with each pause. As expected, more backtracking is detected at pauses in TFIIS KO cells (Figures 2D and S3B). We also determine that persistent backtracking signal can be detected within 50bp of 41% and 84% of WT and TFIIS KO pol II pauses respectively.

However, when comparing LORAX-seq signal to mNET-seq signal at each pause, we find that there is no correlation (Figures 2E and S3C). This analysis indicates that more backtracking is detected in TFIIS KO cells and that while persistent backtracking is mainly found where pol II pauses, pause strength does not determine the potential for persistent backtracking.

DNA sequence has previously been established to affect pausing, due to interactions between elongating pol II and nucleic acids. Specifically, promoter-proximal pausing of pol II is observed to occur at G/C rich regions^8,24,26,27^. We also observe an enrichment for guanine at both WT and TFIIS KO pauses across the genome (Figures 2F and S3D). These results differ from what is observed in yeast, where the consensus sequence around *Δdst1* pauses changes due to large global shifts in pausing (Churchman and Weissman, 2011). The similarity in the consensus sequence of pauses from WT and TFIIS KO cells reiterates the finding that their mNET-seq profiles are highly similar in mammalian cells (Figures 1D-E).

At positions with persistent backtracking, we observe an enrichment for thymine (Figures 2G, S3E-H). The structure of backtracked polymerase indicates that backtracked RNA binds a conserved site in the pol II pore and funnel that preferentially accommodates pyrimidines^2^. *Δdst1* pauses also share a preference for thymine at transiently backtracked positions^20^. This shows that persistent backtracking exhibits a sequence bias unique from pausing. Together, these results indicate that while persistent backtracking occurs at positions where pol II pauses, pausing does not always result in persistent backtracking.

### The footprint of pol II is found proximal to pauses with persistent backtracking

Our data suggests that pol II can be found at positions in the genome proximal to the position of pausing determined by nascent transcript sequencing. To test this, we adapted nascent elongation transcript sequencing to include RNase I footprinting, similar to recent work done in prokaryotes^28,29^. We modified the protocol for use in mammalian cells and importantly, introduced a 4-thiouridine crosslinking step prior to cell lysis in order to capture transcription complexes at their precise position *in vivo* and prevent movement of pol II during sample preparation (Figure 3A, Methods). RNase I would be expected to digest backtracked RNA and result in an upstream shift in pol II footprints at pauses associated with persistent backtracking (Figure 3A).

**Figure 3.**
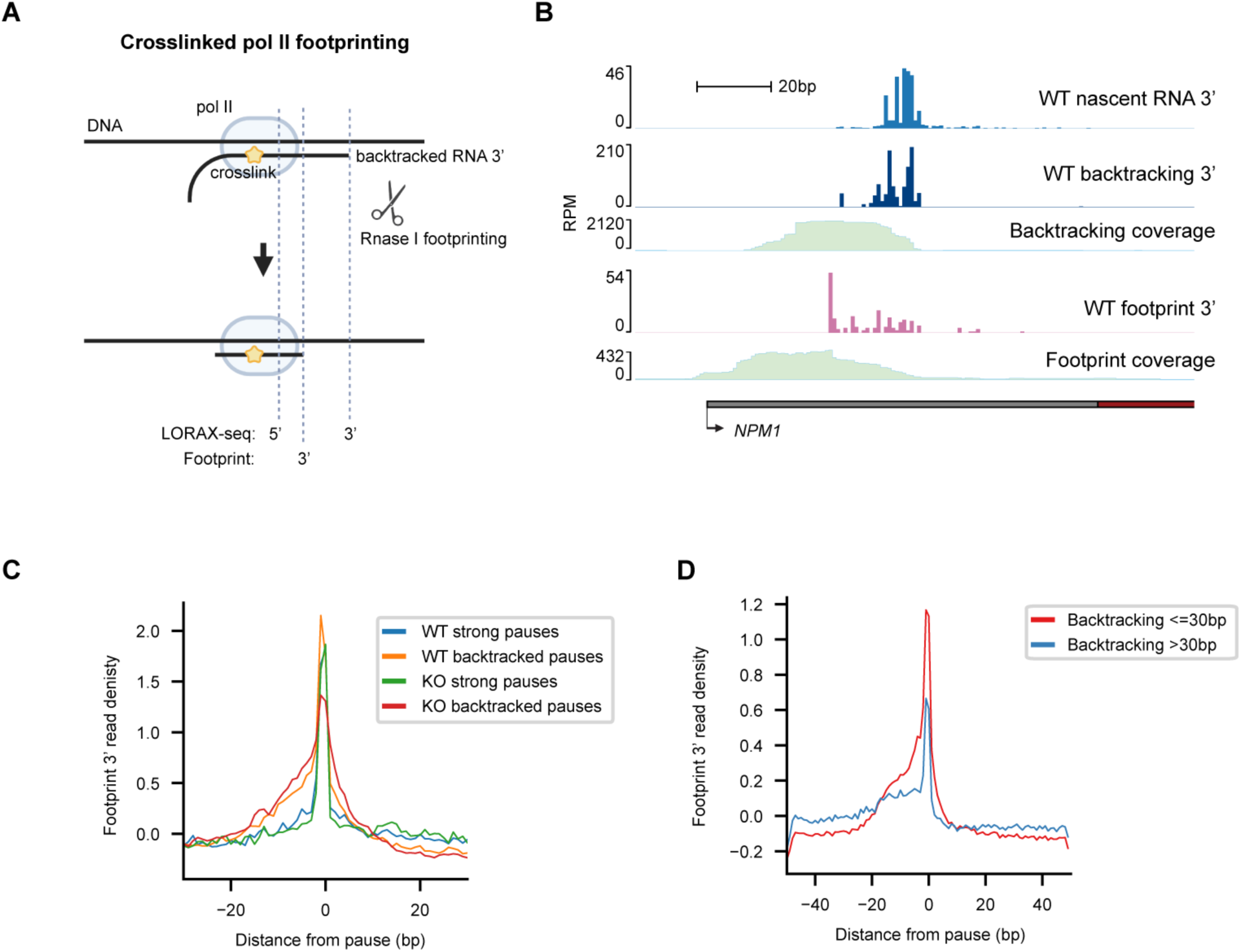
The footprint of pol II is found proximal to pauses with persistent backtracking. (A) Schematic of crosslinked footprinting of pol II. Prior to cell collection, cells were treated with 4-thiouridine and crosslinked. Transcription complexes were collected and Rnase I was used to digest footprints. RNA was then purified and prepared for next generation sequencing. Persistent backtracking of pol II would be expected to shift footprints proximally to the 3’ end detected by LORAX-seq. (B) Representative genome viewer of mNET-seq signal (nascent RNA 3’), LORAX-seq 3’ signal (backtracking 3’), LORAX-seq 5’-3’ signal density showing the extent of backtracking (backtracking coverage), footprint 3’ ends, and footprint full signal density (footprint coverage) at the promoter of NPM1 from WT cells. (C) Meta-analysis of WT and TFIIS KO footprint 3’ ends around n=1,000 pol II pauses with high LORAX-seq signal (backtracked pause) or mNET-seq signal (strong pause) from Figure 2E. Reads within 50 bp of pauses were aggregated, normalized within each window, and averaged. (D) Meta-analysis of TFIIS KO footprint 3’ ends around n=10,000 sites of persistent backtracking less or greater than 30bp in backtracking length. Reads within 50 bp of sites were aggregated, normalized within each window, and averaged. All footprint sequencing experiments were conducted in duplicate for each condition.

Following library preparation and sequencing, the frequency of T to C mutations increased as expected from 4-thiouridine crosslinking (Figure S4A)^30^. In addition, the average footprint size of WT transcription complexes on RNA was around 25nt (Figure S4B). Previous work found pol II transcription complexes to protect either 27 or 60nt of nascent RNA, the latter corresponding with spliceosome associated transcription complexes^31^. We also observed that TFIIS KO resulted in a small increase in footprint length, analogous to the increase observed in cleavage deficient Δ*greAB E. coli* (Figure S4B)^28^. This indicates a larger footprint protected by RNA binding within the backtrack site, as predicted by structure^2^.

At the promoter proximal pause of NPM1 where both mNET-seq nascent RNA and LORAX-seq backtracking signal can be detected, we observe a prominent proximal shift in the footprint of pol II (Figure 3B). To investigate whether this change can be observed at backtracked pauses genome wide, we generated meta-analyses of pol II footprint 3’ ends around pauses from the prior analysis (Figure 2E). As backtracking signal is uncorrelated with pause strength, we compared pauses with high LORAX-seq signal (backtracked pauses) to pauses with high mNET-seq signal (strong pauses). While the footprint 3’ signal is closely associated with the position of strong pauses, it is shifted proximally at backtracked pauses by up to 20 nucleotides (Figures 3C and S4C). In addition, when persistent backtracking events are stratified by backtracking length, the footprint 3’ signal diffuses in the proximal direction around longer backtracking events (Figures 3D and S4D). These results indicate that the nascent RNA 3’ end measured by mNET-seq does not correspond with the position of pol II at persistently backtracked pauses. Furthermore, crosslinked footprints provide additional evidence that stable, long-range backtracking of pol II occurs *in vivo*.

### Genome wide characteristics of persistent backtracking

In a representative comparison of LORAX-seq and mNET-seq signal across the histone gene H1-4, we observe a substantial accumulation of persistent backtracking shortly after the transcription start site (Figure 2B). More signal is found in TFIIS KO cells as previously discussed (Figure 2D); however, the distribution of backtracking across gene bodies is similar in WT cells after normalization (Figure 2B).

To visualize where persistent backtracking occurs genome wide, we produced meta-analyses of LORAX-seq and mNET-seq across transcription start and end sites (TSS and TES) of 10,323 protein coding genes filtered to remove overlapping transcription units. Across the genome, we observe a strong and specific enrichment of persistent backtracking within 250 nucleotides of the TSS, corresponding to the promoter proximal pause^8^ (Figures 4A and S4E-G). Interestingly, we also see persistent backtracking enrichment at the antisense peak, corresponding to bidirectional transcription observed at the TSS^32^ (Figure 4A).

**Figure 4.**
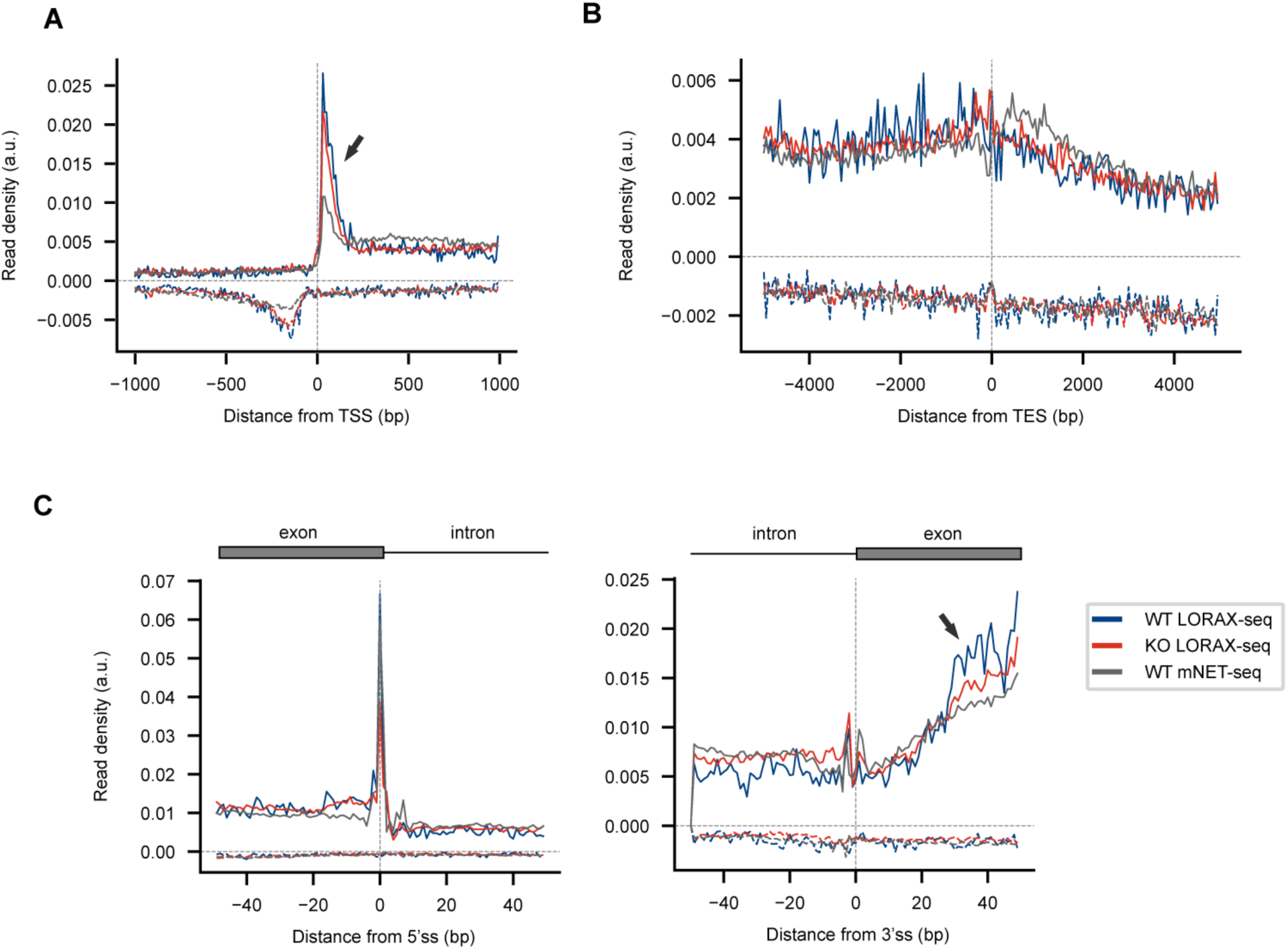
Genome-wide characteristics of persistent backtracking. (A and B) Meta-analysis of persistent backtracking (LORAX-seq) and nascent transcription (mNET-seq) from WT and TFIIS KO cells across the transcription start site (TSS) (A) and transcription end site (TES) (B) of 10,323 protein-coding genes filtered to remove overlapping transcription units. Read density in artificial units (a.u.) is calculated by dividing each window into 200 bins for sense and antisense reads, and then normalizing accumulated reads within each window to be a fraction of one. The sense and antisense strands are presented on the positive and negative y-axes respectively. Arrows denote regions of interest. All LORAX-seq and mNET-seq experiments were conducted in duplicate for each condition. (C) Meta-analysis of LORAX-seq and mNET-seq from TFIIS KO cells across 3’ ends and 5’ ends of splice sites of genes in (A and B).

While we see pausing past the TES with mNET-seq where pol II pauses during termination^22,33^, we did not detect an enrichment for persistent backtracking (Figures 4B and S4H-I). These results specify that persistent backtracking is less prevalent during termination but occurs bidirectionally from the TSS of thousands of human genes and around intron-exon boundaries.

Pol II pausing has been observed after the 3’ splice sites (ss) of intron-exon boundaries to allow for the first catalytic step of co-transcriptional splicing, specifically with pol II phosphorylated at the serine 5 (S5P) CTD position^22^. We observe this phenomenon in our mNET-seq datasets from both WT and KO TFIIS cells (Figure S1L) and detect a corresponding enrichment in LORAX-seq backtracking signal (Figure 4C, arrow; Figure S4J). At exon-intron boundaries, we detect the 5’ss cleavage intermediate as previously described (Figure 4C, left; Figure S4J)^31,34^. Previous work has implicated the dominant negative mutant of TFIIS to affect mRNA splicing^10^ and from our data, we infer that persistent backtracking occurs when pol II pauses during co-transcriptional splicing.

### Effects of persistent backtracking on gene expression

To investigate the functional consequences of persistent backtracking, we determined LORAX-seq signal per gene. We found WT and TFIIS KO LORAX-seq signal per gene to be highly correlated (Figure S5A). Next, we measured gene expression in WT and TFIIS KO cells by RNA seq (Figures S5G-I). Interestingly, genes with LORAX-seq signal above the 95^th^ percentile were significantly downregulated in TFIIS KO cells (Figures 5A-B and S5B-C, Table S1). This suggests that when unresolved, persistent backtracking significantly lowers gene expression.

**Figure 5.**
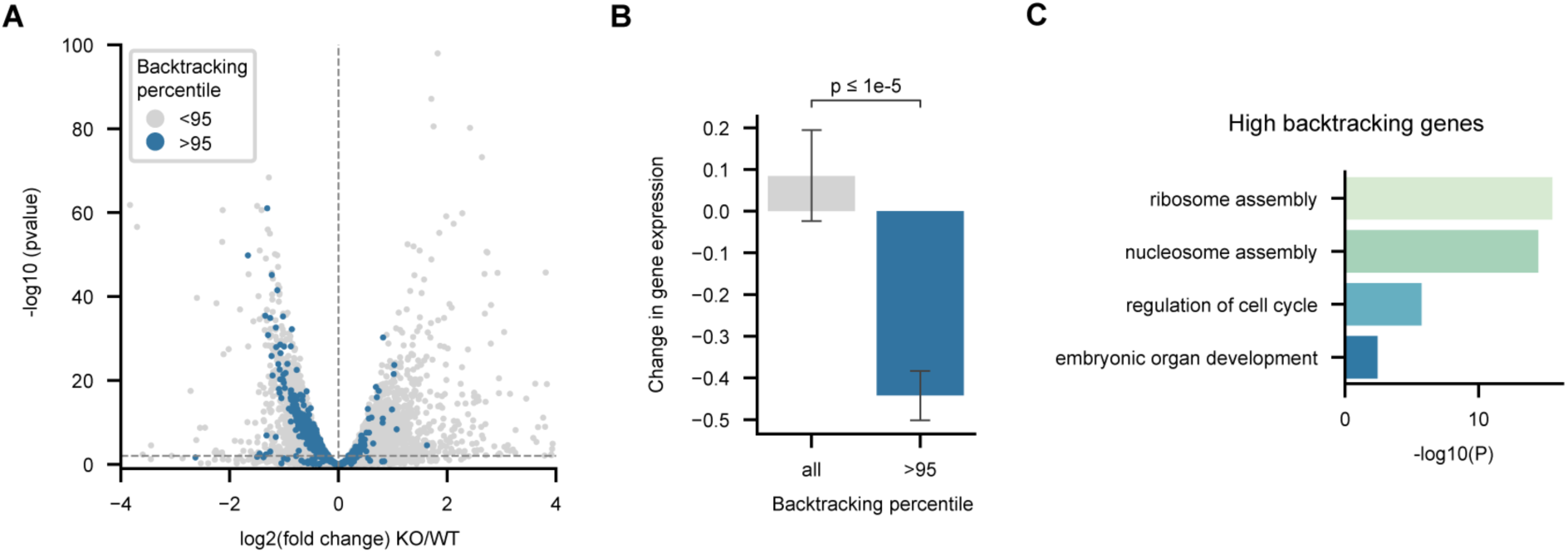
Effects of persistent backtracking on gene expression. (A) Volcano plot of differentially expressed genes between WT and TFIIS KO cells as determined by RNA-seq. Genes with persistent backtracking above the 95^th^ percentile as determined from TFIIS KO LORAX-seq are colored. Horizontal and vertical dotted lines represent p=.01 and a log2 fold change of 0 respectively. The fold change and p values from RNAseq experiments are calculated from 3 biological replicates for each condition. (B) Bar plot of average LFC of genes with high persistent backtracking in (A) compared to all genes. (C) Gene ontology enrichment analysis of 200 genes with high persistent backtracking (>95^th^ percentile).

This also indicates that LORAX-seq is able to identify genes that are likely susceptible to persistent backtracking.

We used gene ontology to identify the function of genes that accumulate persistent backtracking. These genes were found to function in translation, nucleosome assembly, regulation of cell cycle, and development (Figure 5C). We also identified high LORAX-seq signal at heat shock proteins, the prototypical genes for the study of backtracking^15^, and several major transcription factors, such as ATF4 and HOXB9 (Table S1, Figures S5D). Many other genes with high backtracking, particularly those involved in development, were downregulated in TFIIS KO cells (Figures S5H-I).

Highly expressed genes can be transcribed by multiple pol II simultaneously. When a leading polymerase is obstructed, rear-end collisions have been shown to result in extensive backtracking^35^. To test if persistent backtracking is a result of increased gene expression, we compared gene expression to backtracking and observe little correlation (Figure S5E; Pearson R=0.16). Conspicuously, high backtracking genes we find in our analysis such as histone genes do not follow this trend (Figure S5E). Intuitively, LORAX-seq signal per gene moderately correlates with mNET-seq signal, likely because higher pol II occupancy increases the chances for detecting backtracking (Figure S5F). This data indicates that persistent backtracking accumulation at specific genes depends on factors both dependent and independent of gene expression levels.

### Persistent backtracking affects histone gene expression

Histones were among the genes with the highest detected persistent backtracking (Table S1). To investigate the effects of unresolved persistent backtracking at histones, we synchronized cells and measured the expression of histone H2AC20 mRNA by RT-qPCR. While WT cells increase H2AC20 expression 5-fold 4 hours after release from synchronization, TFIIS KO cells increase expression by less than 2-fold compared to timepoint 0 (Figure 6A). Thus, persistent backtracking accumulates on H2AC20 and must be resolved by TFIIS for proper expression during cell division.

**Figure 6.**
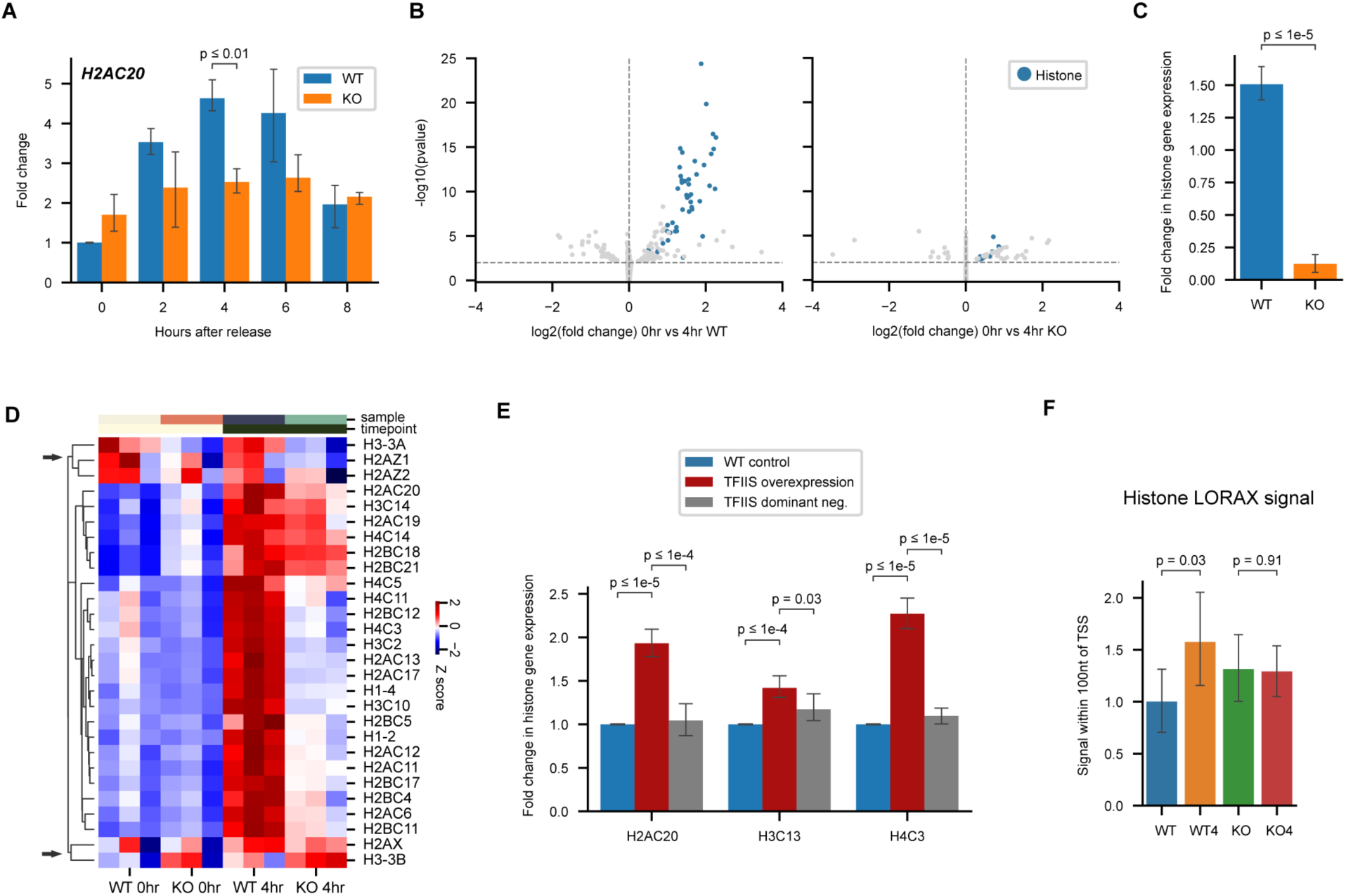
Persistent backtracking coordinates histone gene expression. (A) H2AC20 gene expression after cell synchronization and release as measured by RT-qPCR. Fold changes were normalized to WT at timepoint 0. Bar plots represent the mean of at least 3 biological replicates for each condition, with error bars denoting 95% confidence intervals. (B) Gene expression changes 4 hours after release from synchronization in WT and TFIIS KO cells displayed in RNAseq volcano plots. The log2 fold change (LFC) of cells was determined between 0 and 4 hours after synchronization. Vertical line denotes a LFC of 0 and horizontal line denotes p=.01. Histone genes are colored. The fold change and p values from RNAseq experiments are calculated from 3 biological replicates for each condition. (C) Bar plot displaying average LFC of histone genes in (B) 4 hours after release from synchronization. (D) Heatmap of histone gene expression in WT and TFIIS KO cells 0 and 4 hours after synchronization and release, filtered to remove genes with low expression. Rows are normalized by z score and hierarchically clustered. Arrows denote clusters of replication-independent histone variants (E) Histone gene expression in synchronized WT cells with overexpression of wild-type (overexpression) or cleavage-deficient (dominant neg.) TFIIS, measured by RT-qPCR. Fold changes were normalized to WT control. Bar plots represent the mean of at least 3 biological replicates for each condition, with error bars denoting 95% confidence intervals. (F) LORAX-seq maximum relative signal within 100 nucleotides of the TSS on the sense strand of expressed histone genes from two biological replicates. WT and TFIIS KO cells at 0 and 4 hours after synchronization and release were collected for LORAX-seq analysis. Values were normalized such that the mean WT signal at timepoint 0 is 1.

To test if this also occurs genome wide, we measured gene expression by RNA-seq in WT and TFIIS KO cells 0 and 4 hours after release from synchronization. In WT cells, histone gene expression significantly increases after 4 hours (Figure 6B). Remarkably, no histones were upregulated after 4 hours in TFIIS KO cells (Figure 6C). To examine the variability in impairment of histone expression in KO cells, normalized counts of highly expressed histone genes are plotted in a heatmap and hierarchically clustered (Figure 6D). While the expression of replication-dependent histones is generally impaired in KO cells, there are differences that do not vary with backtracking signal and may result from the timepoint of collection (Figure S6A).

Replication-independent histone variants^36^ form separate clusters and appear to be less affected by TFIIS KO during replication (Figure 6D; arrows).

To investigate if this impairment of histone expression affects replication in TFIIS KO cells, we collected cells after release from synchronization for flow cytometry and propidium iodide staining. Cell cycle analysis indicates that TFIIS KO cells have a prolonged S phase and have difficulty progressing to G2 (Figures S6B-C). Similarly, *Drosophila* cells lacking histones have a prolonged S phase but have been shown to complete DNA replication^37^. These results demonstrate that without TFIIS and resolution of persistent backtracking, cells are unable to properly express histones during S phase.

To investigate if persistent backtracking accumulates at histones prior to their expression during replication, we constructed vectors for overexpression of wild-type and a dominant-negative TFIIS. The dominant-negative differs from wild-type by 2 residues (D190A, E191A) in Domain III that are required for backtracking cleavage and is identical otherwise^2^. Thus, the dominant-negative provides a control for any non-cleavage activities of TFIIS^17^. In synchronized WT and TFIIS KO cells, the overexpression of WT TFIIS significantly increases the expression of three representative histones while the DN mutant TFIIS does not (Figures 6E and S6D).

Overexpression of WT TFIIS likely resolves accumulated backtracking at these loci, increasing histone expression prematurely.

Next, we performed LORAX-seq in synchronized cells to measure how backtracking changes at histone genes. We observe that 4 hours after release from synchronization, LORAX signal increases directly adjacent to the transcription start site in wild-type cells (Figures 6F and S6E). Increased transcription may increase the likelihood of persistent backtracking, which would require TFIIS for resolution. Notably, TSS backtracking in TFIIS KO cells does not change between timepoints (Figures 6F and S6E). Meta-analysis of backtracking signal at all genes remains similar between time points (Figure S6F). Together, these results demonstrate that persistent backtracking accumulates at histone genes and appears to present an obstacle for proper expression during replication.

## Discussion

Backtracking transience and persistence has been predicted *in vitro* in the observation that intrinsic cleavage by pol II can resolve intermediate backtracking lengths while long backtracking can only be resolved with TFIIS^21^. In addition, structural work has indicated that long-range backtracked complexes are stabilized by RNA extruded within the pore and funnel of pol II (secondary channel), impeding the binding of TFIIS^2^, which may explain the persistent nature of long backtracking events.

Nascent transcription sequencing profiles of bacteria, yeast, and mammalian cells lacking the respective cleavage factors have also alluded to these differences. NET-seq in bacteria observes no shift in pausing after deletion of GreA/B, the bacterial cleavage factor, leading to the conclusion that backtracking is rare in *E. coli*^38^. However, later work combining footprinting and NET-seq indicates that backtracking in bacteria occurs frequently without necessarily creating a NET-seq shift in GreA/B mutants (Imashimizu et al., 2015; Sun et al., 2021; Winkelman et al., 2020). NET-seq in yeast is dramatically shifted by up to 18 bp after deletion of Dst1, the yeast cleavage factor^20^, while we and others observe a less pronounced shift in mammalian cells lacking cleavage abilities (Figure 1)^14^. Sequencing of short capped RNA from *Drosophila* has also inferred the existence of long backtracking events at promoter proximal pauses; TFIIS depletion results in an extension of these RNA by up to 40 nucleotides without changing the position of the transcription bubble^40^. These studies demonstrate that depending on the methodology, conclusions about backtracking *in vivo* are starkly different.

We propose to clarify and reconcile these differences by classifying backtracking depending on the ability to observe it in nascent transcription sequencing profiles; transient backtracking can be observed in a state after cleavage but before transcription resumes, while persistent backtracking spends proportionally less time in that state and therefore cannot be observed using nascent transcription sequencing (Figures 1A-B and S1A).

In order to study persistent backtracking, we developed LORAX-seq to directly map the length and precise genomic position where backtracking events begin and end in mammalian cells. Notably, our method uses purified TFIIS to elute backtracked RNA, offering unprecedented sensitivity and specificity for detecting backtracking events. Our data indicates that persistent backtracking mainly originates from positions where pol II pauses and ends 20-40bp upstream (Figure 2C); shorter persistent backtracking events likely also occur but escape detection by sequencing. Transient and persistent events are not mutually exclusive, indicating that transient backtracking might precipitate longer, more persistent events if unresolved (Figure 2C). Additionally, our analysis indicates that while persistent backtracking occurs at positions where pol II pauses, pausing does not always induce persistent backtracking (Figures 2B and 2E).

Long-range and persistent backtracking implies that pol II can be found at positions in the genome proximal to the position of pausing determined by nascent transcript sequencing. To test this, we sequenced the RNA footprint of pol II after crosslinking in order to capture its precise position *in vivo* (Figure 3A). At pauses associated with persistent backtracking RNase I digests backtracked RNA and results in a proximal shift of pol II footprints (Figures 3B and 3C). When backtracking events are stratified by length, the proximal shift is extended around pauses associated with longer backtracking events (Figure 3D). Extensive work with ChIP-seq and derivative methodologies has also observed pol II positioned before the promoter proximal pause^41,42^, where we detect a majority of persistent backtracking (Figure 4). Persistent backtracking may contribute to pol II found in these positions. Furthermore, crosslinked footprints provide additional evidence that pol II can be observed in a persistently backtracked position *in vivo*, divorced from the location of the nascent 3’ end of RNA. This complementary approach provides additional evidence that persistent backtracking occurs *in vivo*.

While we detect more persistent backtracking in TFIIS KO cells (Figure 2D), its distribution across the genome remains consistent (Figure 4). Meta-analysis of LORAX-seq signal indicates that persistent backtracking is ubiquitous but mainly found bidirectionally at promoter-proximal pauses and intron-exon junctions, but less so during termination where transient backtracking can be detected (Figure 4)^14^. Divergent persistent backtracking at the TSS may contribute to the formation of R loops that stimulate antisense transcription^10,43–45^, and TFIIS has been shown to occupy this region by ChIP-seq^14^. In addition, persistent backtracking at intron-exon junctions likely influences the co-transcriptional nature of splicing (Figure 4C)^22,34,46^. This might explain why overexpression of dominant-negative TFIIS causes changes in splicing^10^ despite the absence of transient backtracking at splice junctions^14^. This data shows that persistent and transient backtracking occur at different phases of transcription and that transient backtracking does not always transition into a persistent event.

To explore the functional consequences of persistent backtracking, we stratified genes by LORAX-seq signal and measured their change in expression in TFIIS KO cells. In genes prone to persistent backtracking, gene expression is decreased when backtracking cannot be resolved (Figure 5A and 5B). The high stability of persistent backtracking events appear to act as an obstacle for normal gene expression. We identify the genes most susceptible to persistent backtracking to function in translation, nucleosome assembly, regulation of cell cycle, and development (Figure 5C). Backtracking has previously been implicated in the rapid activation of stress response genes^3,14–16,40^. Our data directly shows that persistent backtracking occurs at these genes *in vivo* in addition to thousands of others previously unknown to be modulated by backtracking and that persistent backtracking accumulates in these genes reliant on factors both dependent and independent of gene expression (Figure 5). Whether the consequences of persistent backtracking are due to hindrance of transcription or premature termination will need to be explored^47^.

Histone genes amass persistent backtracking events most conspicuously, and we provide evidence that the accumulation and resolution of such backtracking is critical for their timely expression. We show that histone genes are not properly expressed in cells unable to resolve backtracking during cell cycle (Figures 6A-D), and that the accumulation of persistent backtracking prevents premature histone expression (Figure 6E-F). We demonstrate that these effects are specifically dependent on the cleavage-stimulation activity of TFIIS (Figure 6E). Pol II pausing has recently been implicated in coordinating cell cycle transition during erythropoiesis^48^. In support, our work provides evidence that backtracking is involved in the timely synthesis of the enormous amounts of histones required for chromatin assembly during replication.

While not a focus of this manuscript, we also found persistent backtracking in genes involved in tissue development (Figures 5 and S5). Intuitively, timely transcription of these genes would be necessary during development, which might explain why TFIIS KO is embryonic lethal in mice^49^. Notably, pol II pausing is implicated in the rapid activation of developmental genes in Drosophila and *C. elegans*^3,8,41,50^. We predict that LORAX-seq in cells undergoing differentiation will provide valuable insights into the role of persistent backtracking in development.

Applying LORAX-seq in different contexts, such as aging or disease, can also increase our understanding of how persistent backtracking affects these processes. It is understood that the heat shock response deteriorates during aging^51^ and interestingly, histone biosynthesis is also defective in aging, resulting in DNA damage, redistribution of epigenetic marks, and ultimately senescence^52^. In both aging and cancer, unresolved persistent backtracking could result in genome instability, chromatin remodeling, and gene dysregulation^10,53–55^. Future work exploring the role of persistent backtracking will broaden our understanding of how it might affect these processes in disease.

## Limitations of the study

This work provides evidence of long ranged, persistent backtracking in mammalian cells and its effects on gene expression using LORAX-seq and crosslinked footprinting of pol II. Notably, the sequencing methods developed in this work have the following limitations: (1) TFIIS treatment of immunoprecipitated transcription complexes requires *in vitro* manipulation, which may not simulate natural conditions. We utilize crosslinked footprinting to provide complementary evidence of persistent backtracking. (2) Nascent transcript sequencing methods, including mNET-seq and sequencing done in this study are relative, providing data on where signal is enriched across the genome within a sample. Future work can incorporate UMIs and spike-ins during library preparation to provide absolute quantification (Gajos et al., 2021). (3) mNET-seq and LORAX-seq are antibody dependent, and only survey captured pol II complexes.

Comparisons between the two should be made using the same antibody. Other limitations of this work include the use of a single human cell line and TFIIS KO, which may cause indirect effects. Future work will need to be done to calibrate LORAX-seq for use in other cell lines and tissues. Finally, our distinction between “transient” and “persistent” backtracking was made based on their detection by mNET-seq and LORAX-seq respectively. Backtracking likely occurs over a spectrum of varying length and persistence.

## Acknowledgments

We thank P. Zappile from the NYU Genome Technology Center, which is partially supported by the Cancer Center Support Grant, P30CA016087, at the Laura and Isaac Perlmutter Cancer Center. We also thank V. Svetlov for help with protein purification and D. Reinberg and G. LeRoy for providing pol II.

## Author contributions

Conceptualization: KBY, AR, EN; Investigation: KBY, CM, TN, IS, VE; Visualization: KBY; Funding acquisition: EN; Supervision: EN; Writing: KBY, EN

## Methods

### Resource Availability

#### Lead contact

Further information and requests for reagents and resources should be directed to and will be fulfilled by the Lead Contact, Dr. Evgeny Nudler.

#### Material availability

Materials generated in this study are available upon request from the Lead Contact with a completed Material Transfer Agreement.

#### Data and code availability

All sequencing data generated are deposited in NCBI’s Gene Expression Omnibus (GEO) database under the accession number GSE235830. This paper does not report original code. Any additional information required to reanalyze the data reported in this paper is available from the lead contact upon request.

### Experimental Model and Subject Details

#### Cell lines and culture conditions

HEK293T/17 cells (ATCC) were maintained in DMEM (ThermoFisher) supplemented with 10% fetal bovine serum (FBS) at 37 °C, 5% CO2 for all experiments.

### Method Details

#### Generation of cell line

CRISPR-Cas9 guides were designed and cloned into the pSpCas9(BB)-2A-Puro (PX459) V2.0 plasmid^56^ to knockout TCEA1 (human TFIIS) in HEK293T/17 cells. Guide RNA sequence is CACCGCCTTGCAGTCCACAAGAATGTTT. Cloning was done using a Gibson Assembly kit (NEB) and NEB5a competent cells (NEB) with oligos ordered from Integrated DNA Technologies (IDT).

#### Overexpression of wild-type and dominant-negative TFIIS

Wild-type and cleavage deficient (dominant negative) TFIIS with mutations D190A and E191A were cloned into the pCDNA3.1 (Invitrogen) plasmid for mammalian expression. Plasmid DNA was prepared using a maxiprep kit (Zymo Research) and transient transfection was achieved by using a 3:1 ratio of PEI (Polysciences, Inc.) to DNA in Opti-MEM (Invitrogen).

#### Western blot

For western blot assays, 1E6 cells were collected and fractionated using the NE-PER nuclear and cytoplasmic extraction kit (ThermoFisher). Antibodies used: Rpb1 CTD (4H8) mouse mAB (#2629; Cell Signaling), TCEA1 (B-6; Santa Cruz), B-actin (C4; Santa Cruz).

#### Purification of human TFIIS

TFIIS was expressed and purified as described previously^6^ with alterations. TFIIS ORF was cloned into pET45b(+) (Millipore Sigma), resulting vector was transformed into Rosetta-gami 2(DE3) cells (Millipore Sigma), overexpression was induced overnight at 30C using Dual Media (Zymo Research). Cells were lysed by sonication in lysis buffer (50 mM Hepes (pH 8.0), 500 mM NaCl, 5% glycerol, 1mM TCEP), supplemented with Lysonase (Millipore Sigma) and ProBlock Gold Bacterial Protease Inhibitor Cocktail (Gold Biotechnology). Lysate was cleared by centrifugation, 29500 g, 60 min, and loaded on 5 ml HisTrap FF Crude column (Cytiva). The column was washed by lysis buffer, supplemented with 10 mM recrystallized imidazole (Gold Biotechnology), and eluted by gradient of 10 to 500 mM of the same in lysis buffer. Peak fractions containing TFIIS were loaded on Superose Increase 6 10/300 GL column equilibrated with 50 mM Hepes (pH 8.0), 1 M NaCl, 5% glycerol, 1 mM TCEP, and eluted in the same. Peak fraction containing TFIIS was dialyzed against TFIIS storage buffer (10 mM Tris-HCl pH 7.5, 500 mM NaCl, 50% glycerol, 5 uM TCEP).

#### TFIIS cleavage of assembled transcription complexes *in vitro*

RNA and DNA oligos were purchased from IDT with PAGE purification. Human pol II was provided by D. Reinberg. RNA-DNA scaffold was assembled by combining 10uL 100uM 5’ TCG AGG TAG CTT GAC GCC TGG TCA AA DNA oligo with 10uL 100uM 5’ -/56-FAM/ rUrGrC rArUrA rArArG rArCrC rArGrG rC RNA oligo in 100uL annealing buffer (12% glycerol; 20mM Tris HCl pH 8.0; 5mM MgCl2; 40mM KCl)^57^. Sample was incubated in PCR cycler at 95 °C for 2 minutes and temperature was dropped from 95 °C to 20 °C in 1 sec intervals, increasing by 1 sec at each 1 degree drop. Resulting scaffold was ethanol precipitated and re-dissolved in 100 ul water to 10 uM concentration. Pol II-scaffold complex was assembled in 40 uL transcription buffer (100mM NaCl; 20mM HEPES pH=7.5; 3mM MgCl2; 4% glycerol, 1mM DTT) by mixing equimolar concentrations of the enzyme and DNA-RNA scaffold (0.2 uM each) for 30 minutes at room temperature. Resulting complex was split into four 10 uL aliquots and TFIIS was added to a concentration of 0.2, 0.5 or 1 uM. Control sample was mixed with equal amount of the mock buffer. Samples were incubated 60 minutes at 28 °C before quenching with 10 uL stop buffer (1XTBE, 20mM EDTA; 8M Urea, 0.025% xylenthianol, 0.025% bromophenol blue). Samples were heated for 5 minutes at 100 °C dry bath and loaded on 15% (20x20cm) (19:1) polyacrylamide gel with 7M Urea and TBE (0.6mm thickness). The gel was run 40 minutes at 50W, transferred to a clean glass and the cleavage products were visualized by Typhoon imager at an excitation wavelength of 473 nm and emission wavelength of 520 nm.

#### LORAX-seq

LORAX-seq was performed by isolating transcription complexes from human cells using an adapted mNET-seq^22^ protocol with modifications from the subsequent POINT method^23^. Starting with 40E6 cells, nuclear fractionation was done as described with HLB+N and HLB+NS buffers. Then, nuclei were resuspended in NUN1 and NUN2 with 3% Empigen (Sigma) and the chromatin was washed with PBS and treated with 50ul TURBO DNase (ThermoFisher), 100ul 10x digestion buffer (100mM Tris-HCl pH 7.5, 25mM MgCl2, 1mM CaCl2), and 10ul Rnasin (Promega) in a volume of 1mL for 15 minutes at 37 °C. Next, Pol II is immunoprecipitated with Rpb1 CTD (4H8) mouse mAB (#2629; Cell Signaling) conjugated to Dynabeads M-280 sheep anti-mouse IgG (ThermoFisher) for 1 hour at 4 °C. Isolated transcription complexes are washed with cold NET-2 buffer with 1% Empigen 5 times before being treated with 100ul cleavage buffer (40mM Tris-HCl pH 7.5, 100mM NaCl, 10mM MgCl2), 2.5ul rSAP (NEB), and 1ul RNasin for 10 minutes at 37 °C, shaking. Magnetic beads were then washed with cold NET-2 with 1% Empigen an additional 5 times before being treated with 50ul cleavage buffer, 2.5uM human TFIIS, and 1ul RNasin for 30 minutes at 37 °C, shaking. Trizol (ThermoFisher) and the Direct-zol RNA microprep (Zymo) was used to purify RNA from the magnetic beads. RNA was then prepared for sequencing with the NEBNext Small RNA library Prep kit (NEB). Sequencing libraries were visualized using an Agilent 2200 Tapestation and sequencing was done using the Novaseq 6000 SP300 cartridge (Illumina).

#### mNET-seq and crosslinked footprinting of pol II

mNET-seq was done essentially as described^22^, similar to the above LORAX-seq protocol except with MNase (NEB) instead of DNase for chromatin digestion, and immediate collection of bead bound RNA and transcription complexes in Trizol following immunoprecipitation by Rpb1 CTD (4H8) mouse mAB conjugated to Dynabeads M-280 sheep anti-mouse IgG (ThermoFisher). mNET-seq experiments were completed in biological replicate. In addition, recent work has improved upon mNET-seq by utilizing unique molecular identifiers (UMI)^24^. Although we do not incorporate UMIs, we conducted all following analyses both with and without computational removal of alignment duplicates. For crosslinked pol II RNA footprinting, mNET-seq was modified in the following ways. Prior to the experiment, cells were treated with 100uM 4-thiouridine (Sigma) for 12 hours. Immediately before collection, media was removed and cells were crosslinked at >310nm at .15J/cm^2, similar to PAR-CLIP^30^. Transcription complexes were then isolated and treated with Rnase I (ThermoFisher) in NET-2 buffer for 5 minutes at 37C, washed with NET-2, and digested with proteinase K (ThermoFisher) in elution buffer (50mM Tris-HCl pH 7.5, 10mM EDTA, 1% SDS) for 90 minutes at 55 °C before collection in Trizol.

Purified RNA was treated with polynucleotide kinase (NEB) to prepare 5’ and 3’ ends for library construction as described for LORAX-seq.

#### Cell cycle analysis

Cell cycle synchronization was achieved using double thymidine block. 0.1E6 cells were seeded in a 12 well plate. The next day, media was replaced with normal growth medium supplemented with 2mM thymidine (Sigma) and cultured overnight for 18 hours. Media was replaced with normal growth medium for 8 hours and then replaced again with 2mM thymidine medium for 18 hours. The next day, normal growth medium is added to release cells from synchronization. Cells were collected at the indicated timepoints for staining with propidium iodide to measure DNA content. Samples were collected with a FACSCalibur flow cytometer (BD Biosciences).

#### RNAseq

RNA was collected from cells using Trizol and Direct-zol RNA microprep. Cell synchronization is described above. The NEBNext rRNA depletion kit (NEB) was used to deplete ribosomal RNA. Note that poly-A enrichment would exclude non-polyadenylated histone genes. RNA-seq libraries were then prepared using the NEBNext Ultra II directional RNA library kit (NEB).

#### RT-qPCR experiments

RNA was prepared from synchronized cells using Trizol and Direct-zol RNA microprep. RT-qPCR was done with 100ng of RNA with the Luna RT-qPCR kit (NEB) using primers targeting beta actin as a control (Table S2). For wild-type and cleavage deficient TFIIS overexpression experiments using transient transfection, a single round of thymidine block was used to synchronize cells.

### Quantification and Statistical Analysis

#### RNAseq analysis

Sequencing adapters were first trimmed with CutAdapt^58^ and then aligned to the hg19 reference genome using HISAT2^59^. Salmon^60^ was used to quantify expression and DESeq2^61^ was used to identify differentially expressed genes.

#### LORAX-seq and mNET-seq data analysis

Sequencing adapters were first trimmed with CutAdapt^58^ and then aligned to the hg19 reference genome using HISAT2^59^. Alignment files were converted to bigwig format with CPM normalization using bamCoverage from deepTools2^62^ for genome browser visualization with pyGenomeTracks^63^. For other analyses, alignment files were converted to bed format and annotated using Samtools and BEDtools^64,65^. Scripts were written to deduplicate reads, extract read lengths, produce meta-analysis plots, and run gene ontology analyses. Meta-analysis plots were produced by collecting sense and antisense reads in 200 bins (100 on each strand) for selected genomic regions with detectable signal and normalizing the reads to be a fraction of one. Genomic regions include pol II pauses, splice sites, and transcription start and end sites, which were filtered to include only protein coding genes and remove overlapping regions within 2.5 kilobases. Gene ontology analyses were run using the web tool provided by the Gene Ontology Consortium^66,67^ and PANTHER^68^. mNET-seq pol II pauses were called with an algorithm similar to those used previously^20^. Pauses were identified in positions where there are at least 5 mapped reads and the signal is > 3 standard deviations above the mean signal in the surrounding 200 nucleotides. Consensus sequences were determined and plotted using Logomaker^69^. ’Backtracking index’ was calculated by determining the difference in read density upstream and downstream of each pause in WT and KO mNET-seq as previously described^14^.

